## Supplemental Figures for "Causal Haplotype Block Identification in Plant Genome-Wide Association Studies"

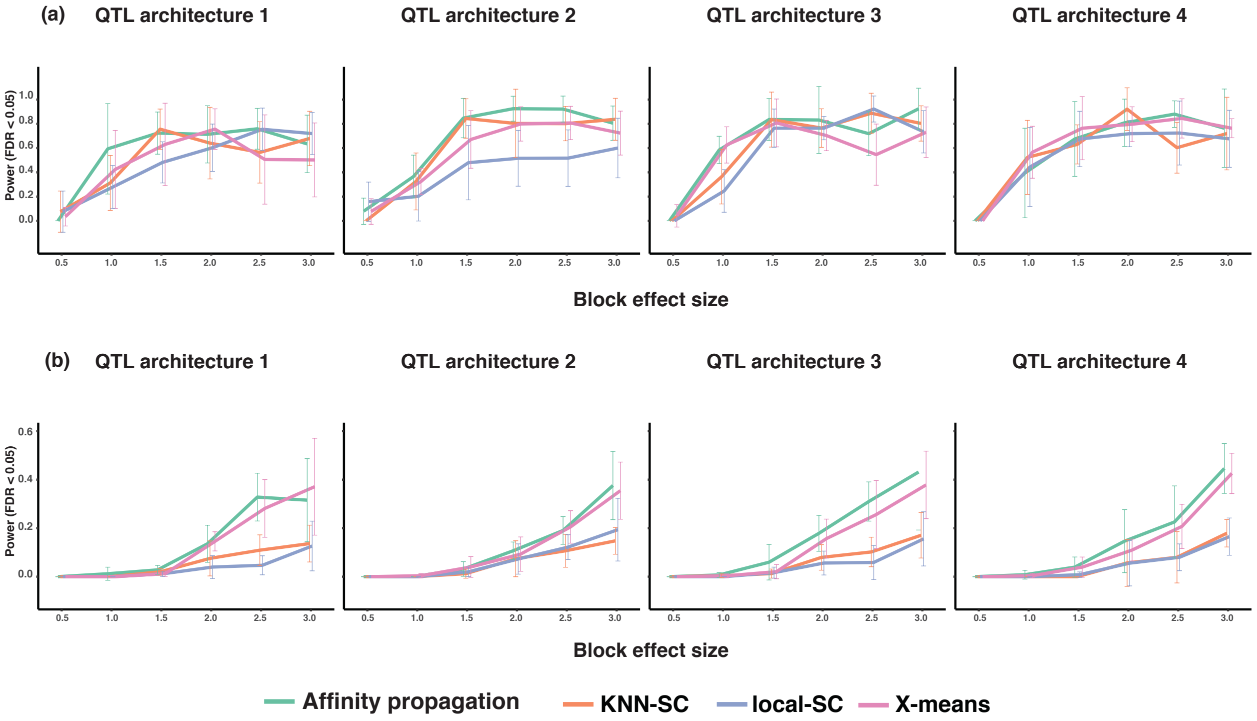


Supplemental Figure 1. Mapping power comparison of different haplotype clustering algorithms.

(a) Mapping power (FDR < 0.05) of affinity propagation, KNN-spectral clustering, local-spectral clustering and X-means in the low polygenicity simulation

(b) Mapping power (FDR < 0.05) of affinity propagation, KNN-spectral clustering, local-spectral clustering and X-means in the high polygenicity simulation

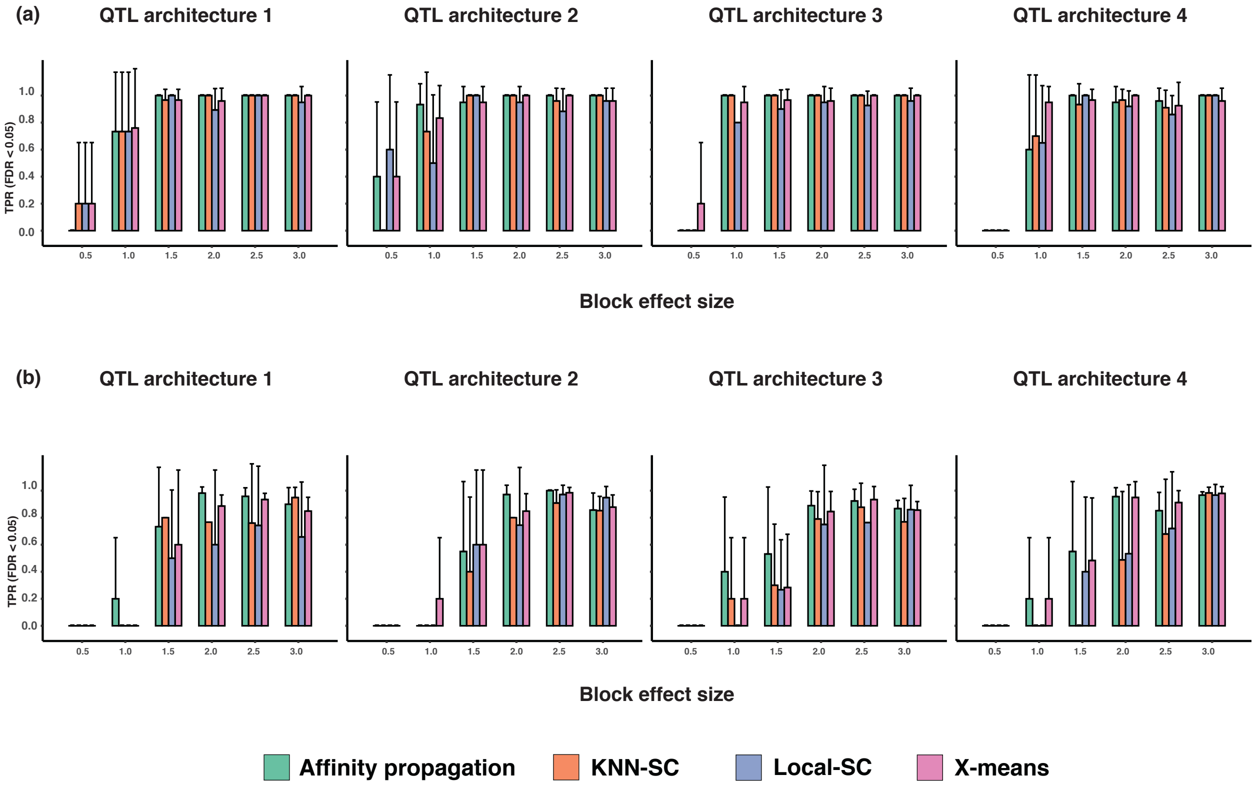


Supplemental Figure 2. True positive rate (TPR) comparison of different haplotype clustering algorithms.

(a) TPR (FDR < 0.05) of affinity propagation, KNN-spectral clustering, local-spectral clustering and X-means in the low polygenicity simulation

(b) TPR (FDR < 0.05) of affinity propagation, KNN-spectral clustering, local-spectral clustering and X-means in the high polygenicity simulation

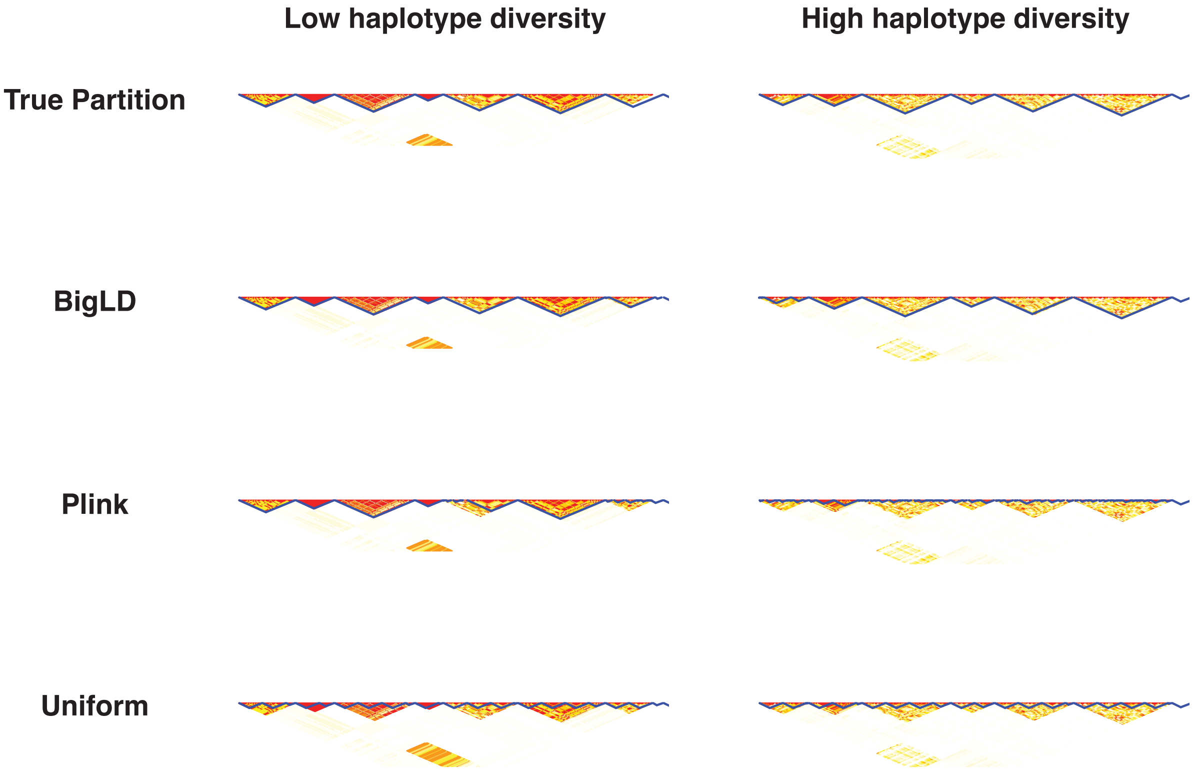


Supplemental Figure 3. Comparison of Block partition algorithms on simulated datasets. Each block partition algorithm was tested on low haplotype diversity and high haplotype diversity simulations. The redness indicates the strength of LD between SNP pairs, and the blue line indicates the block partition generated by the method.

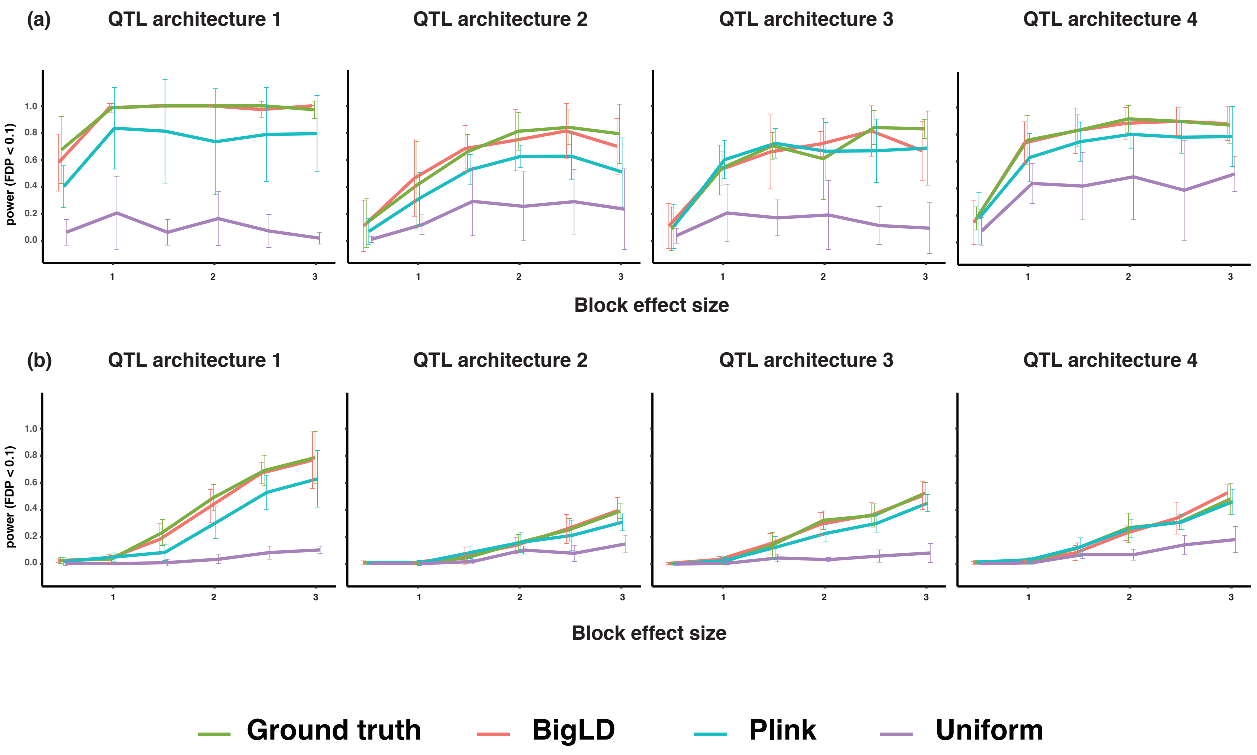


Supplemental Figure 4. Mapping power comparison of different block partition algorithms in the low polygenicity simulations. The x-axis indicates the per-locus heritability.

(b). Mapping power comparison (FDR < 0.05) of block partition algorithms in the low haplotype diversity and low polygenicity simulations.

(c). Mapping power comparison (FDR < 0.05) of block partition algorithms in the high haplotype diversity and low polygenicity simulations.

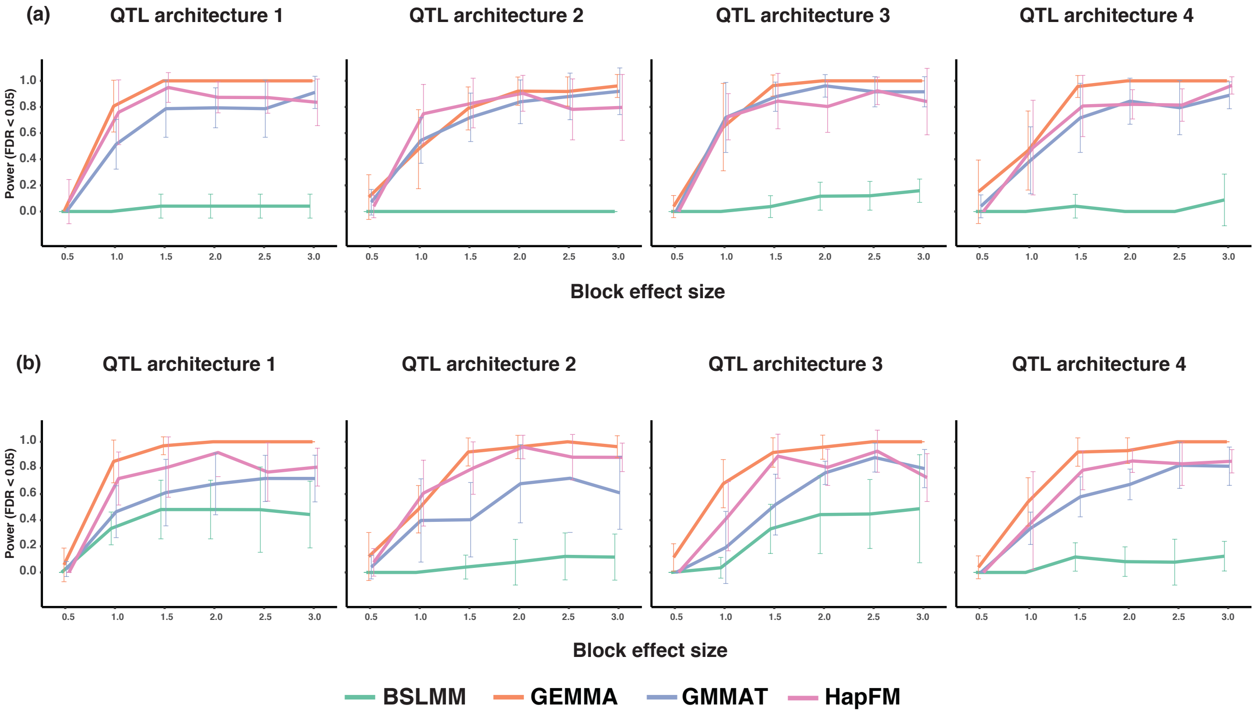


Supplemental Figure 5. Mapping power comparison of different GWAS algorithms in the low polygenicity simulations. The x-axis indicates the per-locus heritability.

(a). Mapping power comparison (FDR < 0.05) of different GWAS algorithms in the low haplotype diversity and low polygenicity simulations.

(b). Mapping power comparison (FDR < 0.05) of different GWAS algorithms in the high haplotype diversity and low polygenicity simulations.

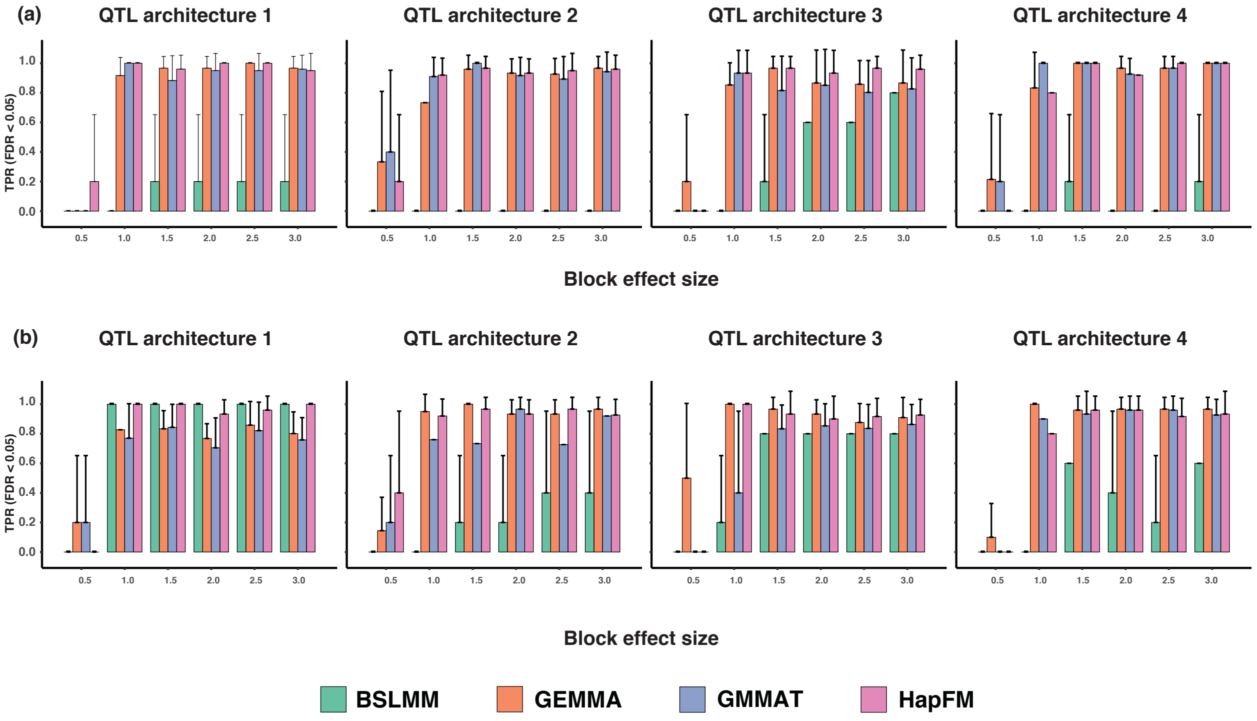


Supplemental Figure 6. True positive rate different GWAS algorithms in the low polygenicity simulations. The x-axis indicates the per-locus heritability.

(a). True positive rate (FDR < 0.05) of different GWAS algorithms in the low haplotype diversity and low polygenicity simulations.

(b). True positive rate (FDR < 0.05) of different GWAS algorithms in the high haplotype diversity and low polygenicity simulations.

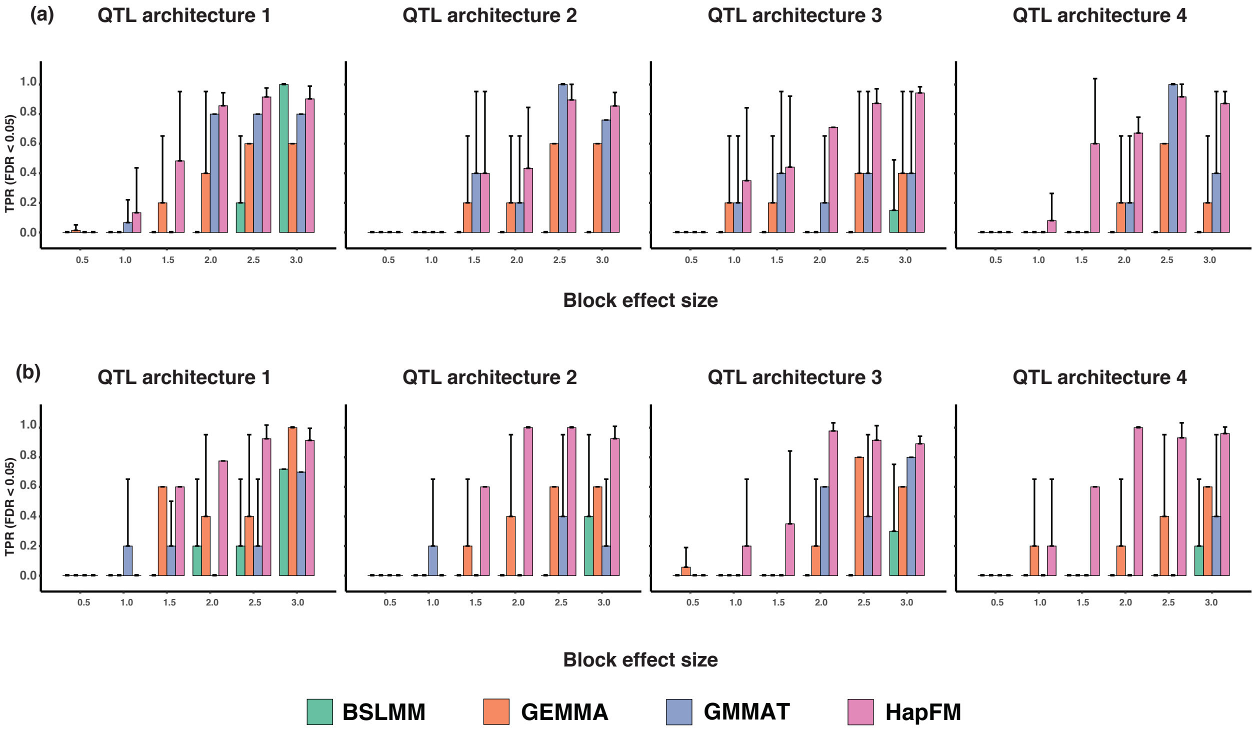


Supplemental Figure 6. True positive rate different GWAS algorithms in the high polygenicity simulations. The x-axis indicates the per-locus heritability.

(a). True positive rate (FDR < 0.05) of different GWAS algorithms in the low haplotype diversity and low polygenicity simulations.

(b). True positive rate (FDR < 0.05) of different GWAS algorithms in the high haplotype diversity and low polygenicity simulations.
